# Induced systemic resistance impacts the phyllosphere microbiome through plant-microbe-microbe interactions

**DOI:** 10.1101/2021.01.13.426583

**Authors:** Anna Sommer, Marion Wenig, Claudia Knappe, Susanne Kublik, Bärbel Fösel, Michael Schloter, A. Corina Vlot

## Abstract

Both above- and below-ground parts of plants are constantly confronted with microbes, which are main drivers for the development of plant-microbe interactions. Plant growth-promoting rhizobacteria enhance the immunity of above-ground tissues, which is known as induced systemic resistance (ISR). We show here that ISR also influences the leaf microbiome. We compared ISR triggered by the model strain *Pseudomonas simiae* WCS417r (WCS417) to that triggered by *Bacillus thuringiensis israelensis* (*Bti*) in *Arabidopsis thaliana*. In contrast to earlier findings, immunity elicited by both strains depended on salicylic acid. Both strains further relied on MYC2 for signal transduction in the plant, while WCS417-elicited ISR additionally depended on SAR-associated metabolites, including pipecolic acid. A metabarcoding approach applied to the leaf microbiome revealed a significant ISR-associated enrichment of amplicon sequence variants with predicted plant growth-promoting properties. WCS417 caused a particularly dramatic shift in the leaf microbiota with more than 50% of amplicon reads representing two bacterial species: WCS417 and *Flavobacterium* sp.. Co-inoculation experiments using WCS417 and *At-*LSPHERE *Flavobacterium* sp. Leaf82, suggest that the proliferation of these bacteria is influenced by both microbial and plant-derived factors. Together, our data connect systemic immunity with leaf microbiome dynamics and highlight the importance of plant-microbe-microbe interactions for plant health.

## Introduction

The functional traits introduced by the plant-associated microbiome are essential for plant growth and fitness and include nutrient acquisition as well as improved responses of the plant towards abiotic and biotic stressors (Berg, 2009; Schlaeppi & Bulgarelli, 2015). Some microbes are able to activate plant defence mechanisms, including systemic acquired resistance (SAR) and induced systemic resistance (ISR). While SAR is induced in systemic tissues of plants undergoing a local pathogen infection, ISR takes effect in aerial tissues of plants interacting with beneficial microbes in the rhizosphere (Vlot et al., 2020).

The molecular mechanisms of SAR are well-researched. SAR depends on two distinct but interwoven signalling pathways, one depending on salicylic acid (SA), the other on pipecolic acid (Pip) (Vlot et al., 2020). SA levels rise both locally and systemically after pathogen infection. This is driven by the enzymes ISOCHORISMATE SYNTHASE 1 (ICS1, also known as SID2) followed by the amidotransferase AvrPphB SUSCEPTIBLE3 (PBS3) (Rekhter et al., 2019; Vlot, Dempsey, & Klessig, 2009; Wildermuth, Dewdney, Wu, & Ausubel, 2001). Elevated SA levels lead to enhanced resistance through the action of downstream signalling intermediates, including the proposed SA receptors NON-EXPRESSOR OF PATHOGENESIS-RELATED PROTEINS1 (NPR1) and its paralogs NPR3 and 4 (Cao, Glazebrook, Clarke, Volko, & Dong, 1997; Y. Ding et al., 2018; Fu et al., 2012; Liu et al., 2020). In parallel, the non-proteinogenic amino acid Pip is synthesized in two steps by AGD2-like Defence Response Protein1 (ALD1) and SAR-DEFICIENT 4 (SARD4) and then converted to its presumed bioactive form *N*-hydroxy-pipecolic acid (NHP) (Chen et al., 2018; P. Ding et al., 2016; Hartmann et al., 2017; Hartmann et al., 2018; Navarova, Bernsdorff, Doring, & Zeier, 2012). Notably, SA and Pip are believed to fortify each other’s accumulation in a positive feedback loop, which depends on shared transcription (co-)factors, including NPR1 (Y. Kim, Gilmour, Chao, Park, & Thomashow, 2020; Sun et al., 2020).

The long-distance signal, which mediates the communication between local infected and systemic tissues and ultimately triggers the establishment of SAR, appears to be composed of multiple signalling intermediates, including SA, Pip, and/or NHP (reviewed in (Vlot et al., 2020). Additionally, volatile signals such as the monoterpenes camphene and α- and β-pinene are essential for SAR and propagate systemic immunity in SAR-induced as well as neighbouring plants (Riedlmeier et al., 2017; Wenig et al., 2019). GERANYL GERANYL DIPHOSPHATE SYNTHASE 12 (GGPPS12) is a key enzyme in the production of volatile monoterpenes in *Arabidopsis thaliana*. Mutations in this gene reduce monoterpene emissions and the capacity of the volatile emissions of these plants to support SAR (Riedlmeier et al., 2017). Perception of monoterpenes in SAR depends on the downstream SAR signalling intermediate LEGUME LECTIN-LIKE PROTEIN 1 (LLP1) (Breitenbach et al., 2014; Wenig et al., 2019).

ISR is elicited by plant growth-promoting bacteria or fungi in the rhizosphere (PGPR/PGPF), including, for example, several *Pseudomonas, Bacillus*, and *Trichoderma* strains (Pieterse et al., 2014; Vlot et al., 2020). In contrast to SAR, which provides protection against (hemi-) biotrophic pathogens, ISR protects above-ground tissues against both necrotrophic and (hemi-) biotrophic pathogens (Pieterse, van Wees, Hoffland, van Pelt, & van Loon, 1996; Ton, Van Pelt, Van Loon, & Pieterse, 2002; Van der Ent et al., 2008; Waller et al., 2005). The best-characterized ISR system to date is that induced in *Arabidopsis thaliana* upon interaction of the roots with *Pseudomonas simiae* WCS417r (Pieterse et al., 1996). The exact mechanism by which the presence of the microbes is perceived at the roots and relayed to the whole plant is not known at this point. The traditional idea is that ISR signals are propagated in the plant via jasmonic acid (JA)- and ethylene (ET)- dependent signalling (Pieterse et al., 1996; Pieterse et al., 1998; Pozo, Van Der Ent, Van Loon, & Pieterse, 2008). However, evidence is accumulating that there is no uniform ISR response to all PGPRs. Instead, there seem to be differing responses, depending on the eliciting microbial strains, involving JA/ET signalling as well as SA signalling pathways (Kojima, Hossain, Kubota, & Hyakumachi, 2013; Martínez-Medina et al., 2013; Nie et al., 2017; Niu et al., 2011; van de Mortel et al., 2012; Wu et al., 2018). These different responses are believed to enable the plant to react in a directed manner dependent on the lifestyle of the attacking pathogen (Nguyen et al., 2020). Signal propagation to the aerial tissues of the plant leads to so-called priming. During priming, full defence responses are not immediately activated. Rather, the plant raises a stronger and faster immune response after pathogenic challenge as compared to unprimed plants (U. Conrath, G. J. M. Beckers, C. J. G. Langenbach, & M. R. Jaskiewicz, 2015; Martinez-Medina et al., 2016; Mauch-Mani, Baccelli, Luna, & Flors, 2017).

The plant immune system influences the propagation of pathogens, but also that of non-pathogenic commensal or plant growth-promoting microbes, which are associated with the plant and together make up the plant microbiota (Teixeira, Colaianni, Fitzpatrick, & Dangl, 2019). Local interactions of plant organs with pathogens can trigger long-distance signalling, for example from leaves to roots, and mediate changes in root exudates that influence the composition of the rhizosphere microbiota (Berendsen et al., 2018; Rudrappa, Czymmek, Pare, & Bais, 2008; Yu, Pieterse, Bakker, & Berendsen, 2019). Similar changes in the plant immune status are associated with the dynamics of the phyllosphere microbiome (Chaudhry et al., 2020). Certain commensal bacteria from the phyllosphere, in turn, have been shown to enhance, for example, SA-associated immunity (Vogel, Bodenhausen, Gruissem, & Vorholt, 2016). It thus seems conceivable that the plant immune system can modulate the phyllosphere microbiome, allowing the plant to ‘exploit’ beneficial properties of microbes to promote plant fitness.

In this study, we show that plant-microbe interactions in the rhizosphere influence the composition of the above-ground phyllosphere microbiome. We combine induced resistance assays in different *A. thaliana* genotypes with a molecular barcoding approach based on sequencing of amplified fragments of the 16S rRNA gene to assess the phyllosphere microbiome. The use of two different ISR inducers, *P. simiae* WCS417r and *Bacillus thuringiensis* var. *israelensis*, allows us to differentiate between local and systemic responses. Importantly, the data suggest that ISR-induced responses of the plant microbiome are influenced by interconnected microbe-microbe and microbe-plant interactions, which in response to *P. simiae* WCS417r reduce species diversity and thus presumably the stability of the leaf microbiome. Our results thus reveal a possible trade-off of ISR-based plant protection strategies and highlight the importance of tri-partite plant-microbe-microbe interactions for plant health.

## Methods

### Plant material and growth conditions

In this study, *A. thaliana* ecotype Columbia-0 (Col-0) was used for all experiments. The mutants *llp1-1, ggpps12, ald1, npr1-1, sid2*, and *jasmonate-insensitive 1* (*jin1*) were previously described (Berger, Bell, & Mullet, 1996; Breitenbach et al., 2014; Cao et al., 1997; Riedlmeier et al., 2017; Song, Lu, McDowell, & Greenberg, 2004; Wenig et al., 2019; Wildermuth et al., 2001). All plants were grown from synchronised seeds. Plants were grown on normal potting soil (“Floradur® B Seed” (Floragard GmbH, Oldenburg, Germany) mixed with silica sand (grain size 0,6-1,2mm) at a ratio of 5:1. For ISR experiments seeds were surface-sterilized with 75% ethanol twice for 4 minutes and grown on ½ Murashige and Skoog medium for 10 days (d) prior to treatment and transfer to soil. Plants were grown in a 10 hour (h) day light regiment and a light intensity of 100µmol m^-2^ s^-1^ photosynthetically active photon flux density at 22°C during light periods and 18°C during dark periods. Relative humidity was kept at >70%.

### ISR elicitors, Pathogens and Treatments

For elicitation of ISR, two different bacterial strains were used: *Pseudomonas simiae* WCS417r (Pieterse et al., 1996) and *Bacillus thuringiensis* var. *israelensis* (Goldberg, 1977). For ISR treatment, bacteria were grown on NB (Carl Roth, Karlsruhe, Germany) plates for 24 h and suspended in 10mM MgCl_2_ to a final concentration of 10^8^ colony forming units (cfu) mL^-1^, assuming that an OD_600_ =1 corresponds to 10^8^ cfu mL^-1^. To induce ISR, the roots of 10-day-old seedlings were placed in wells of 96-well plates containing one of the bacterial suspensions or a sterile 10mM MgCl_2_ control solution, each supplemented with 0.01% Tween-20 (v:v). After 1 h of incubation, the seedlings were placed in pots with soil and grown to an age of 34 d. On the 34^th^ day after sowing, the leaves of the plants were either harvested for further analysis or inoculated with 10^5^ cfu mL^-1^ of *Pseudomonas syringae* pathovar *tomato* (*Pst*), which was maintained and used for infections as previously described (Wenig et al., 2019). To determine bacterial growth in the plants, *Pst* titers were determined 4 days post-inoculation (dpi). To this end, leaf discs punched out of the infected leaves were incubated in 10mM MgCl_2_ + 0,01% Silwet (v:v) for 1 h at 600 rpm. The resulting bacterial suspensions were serially diluted in steps of 10x. 20µl per dilution were plated on NYGA agar plates (Wenig et al., 2019) and incubated for 2 d at room temperature. Bacterial titers were calculated based on the number of bacterial colonies.

Leaf inoculations were performed using 4-5-week-old plants. *Flavobacterium sp*. was obtained as strain Leaf82 from the *At*-LSPHERE synthetic community (Bai et al., 2015) and maintained on NB medium. Syringe infiltration was performed using 10^5^ cfu mL^-1^ of bacteria in 10 mM MgCl_2_. Spray inoculation was performed using 10^8^ cfu mL^-1^ of bacteria in 10 mM MgCl_2_ supplemented with 0.01% Tween-20 (v:v). *In planta* bacterial titers were determined as described above by counting plate-grown bacterial colonies derived from inoculated leaves. The colonies of WCS417 and Leaf82 were distinguished based on colour differences.

SAR was induced in 4-5-week-old plants as previously described (Wenig et al., 2019) except that WCS417 or Bti were used for the primary inoculation of the first two true leaves of the plants by syringe infiltration of 10^6^ cfu mL^-1^ of bacteria in sterile 10 mM MgCl_2_. 10^6^ cfu mL^-1^ of *Pst* carrying the effector *AvrRpm1* was used as the positive control and 10 mM MgCl_2_ as the negative control treatment (Wenig et al., 2019). Three d later, the establishment of SAR was tested by a secondary infection of the third and fourth true leaf of the plants with 10^5^ cfu mL^-1^ of *Pst. Pst* titers were determined at 4 dpi as described above.

### RNA extraction and RT-qPCR analysis

RNA was isolated with Tri-Reagent (Sigma-Aldrich, St. Louis, USA) according to the manufacturer’s instructions. cDNA was generated with SuperscriptII reverse transcriptase (Invitrogen, Carlsbad, USA). Quantitative PCR (qPCR) was performed using the Sensimix SYBR low-rox kit (Bioline, Memphis, USA) on a 7500 real-time PCR system (Applied Biosystems, Foster City, USA). Primers that were used for qPCR are listed in Supplementary Table S1. Transcript accumulation of target genes was analyzed using Relative Quantification with the 7500 Fast System Software 1.3.1.

### DNA-Isolation, PCR and Amplicon Sequencing

100-200ng of leaf material was freeze-dried for 24 h at -40°C and 0.12mbar (Alpha 2-4 LD Plus, Martin Christ Gefriertrocknungsanlagen, Osterode, Germany). DNA isolation was performed utilizing the FastPrep Soil Kit (MPbio) according to manufacturer’s instructions after an additional step of leaf grinding using a tissue lyser (Retsch, Haan, Germany) and glass beads (1mm diameter) at 25Hz for two minutes. Following DNA extraction, the variable regions V5-V7 of the bacterial 16S rRNA gene were amplified by PCR (NEBnext High Fidelity 2x Master Mix, New England Biolabs, Ipswich, MA, USA) using 10 ng of DNA per reaction and the primers 799F and 1193R from (Bulgarelli et al., 2012; Chelius & Triplett, 2001).

Three independent PCR reactions were performed per DNA sample using the following conditions: 98°C for 30 s, 98°C for 10 s, 58°C for 20 s, 72°C for 20 s, 72°C for 2 m. Steps 2-4 were repeated 25 times. The resulting PCR amplicons were subjected to gel electrophoresis to separate amplicons derived from bacteria and chloroplasts, since chloroplast yield longer amplicons than bacterial DNA. The DNA amplicons derived from the bacterial 16S rRNA gene were extracted from the gels using the QIAquick Gel Extraction Kit (Qiagen, Hilden, Germany). After determination of the DNA concentration of each amplicon (nanodrop, Implen, Munich, Germany), the 16S rRNA gene amplicons from 3 replicates per sample were pooled at equimolar amounts. The fragment sizes and concentrations of the pooled samples were determined on a Fragment analyzer 5200 using the DNF-473-Standard Sensitivity NGS Fragment Analysis Kit (Agilent, Santa Clara, CA, USA). The indexing PCR was performed under the following conditions: 98°C for 10 s, 55°C for 30 s and 72°C for 30 s and final extension at 72°C for 5 min. Each PCR reaction contained 1x NEBNext High Fidelity Mastermix, 10 ng of template DNA and index primer 1 (N7xx) and index primer 2 (N5xx) of Nextera XT Index Kit v2 Set A (Illumina, San Diego, CA, USA) according to the manufacturer’s instruction. All samples were purified using MagSi NGSprep Plus Beads (Steinbrenner, Wiesenbach, Germany). Samples were validated and quantified on a Fragment analyzer 5200 using the DNF-473-Standard Sensitivity NGS Fragment Analysis Kit, diluted and pooled to a final concentration of 4 nM for the sequencing run on Illumina MiSeq using the MiSeq Reagent Kit v3 (600-cycle). Demultiplexing was done using the MiSeq Reporter Software v 2.6. (Illumina).

### Statistical analysis

All statistical analyses were done using R version 3.6.3.(R Development Core Team, 2020). For the analysis of bacterial titers, a Shapiro wilk test for normal distribution showed, that the cfu counts resulting from the infection assays did not follow normal distribution (α=0.05). Therefore, a Kruskal-Wallis test was used to test for significance at α=0.05, followed by a post hoc pairwise Wilcox test with correction for multiple testing using the Benjamini&Hochberg method.

### Amplicon data analysis

Pre-processing of the amplicon data was performed using the package “dada2”, including removal of low-quality reads, merging of reads, chimera removal and taxonomic assignment based on the Silva Seeds v138 database (Callahan BJ, 2016; Yilmaz et al., 2013). Taxonomy assignments were performed based on Amplicon Sequence Variants (ASVs). Phylogenetic trees were fitted based on DECIPHER (Wright, 2016). To control for uniformity of DNA isolation and PCR bias as well as contamination, a commercially available Microbial Community Standard by ZymoBIOMICs was prepared as an additional sample and handled in the same fashion as the other samples after the freeze-drying step.

Prior to analysis of the data, we mined the *Pst* titer reductions triggered by each treatment as compared to the appropriate controls, and excluded data from samples derived from experiments, in which ISR was not significant. Data from the remaining 6-7 replicates per treatment were analysed using the R packages Vegan, Phyloseq, DESeq2, and Phangorn were used (Holmes, 2013; Jari Oksanen, 2019; Love, 2014; Schliep, Potts, Morrison, & Grimm, 2017). Read counts were normalized to the read count per sample. Alpha diversity was calculated using the Shannon’s- as well the Simpson’s index (Phyloseq). Nonmetric Multidimensional Scaling (NMDS) was plotted after calculation of Unifrac-distances (Phyloseq). Based on these analyses and Grubbs outlier tests (*p* < 0.05), we excluded the data from one sample per treatment, which were outliers in terms of species-richness in comparison to the other samples of the respective treatments. Differentially abundant ASVs were determined using DESeq2, limiting the analysis to ASVs present in at last three samples.

## Results

### Bacillus thuringiensis var israelensis (Bti) elicits ISR in A. thaliana

*P. simiae* WCS417r, referred to below as WCS417, triggers ISR in *A. thaliana*, reducing the propagation of pathogenic *P. syringae* pathovar *tomato* (*Pst*) in the leaves of the treated plants (Pieterse et al., 1996). Here, we tested if treatment of *A. thaliana* roots with *Bti* has a similar effect. To this end, 10-day-old, sterile-grown seedlings were treated with *Bti* or with WCS417 as a positive control or with sterile 10mM MgCl_2_ as a negative control. The treated seedlings were transferred to soil, and leaves of the resulting plants were inoculated with *Pst*. As expected, treatment of *A. thaliana* roots with WCS417 reduced the growth of the *Pst* inoculum in the leaves as compared to that in control plants, indicating the induction of ISR in response to WCS417 (Fig. 1A). Treatment of seedling roots with *Bti* caused a comparable reduction of *Pst* growth in the leaves (Fig. 1A), indicating that *Bti* induced ISR in *A. thaliana*.

**Figure 1.**
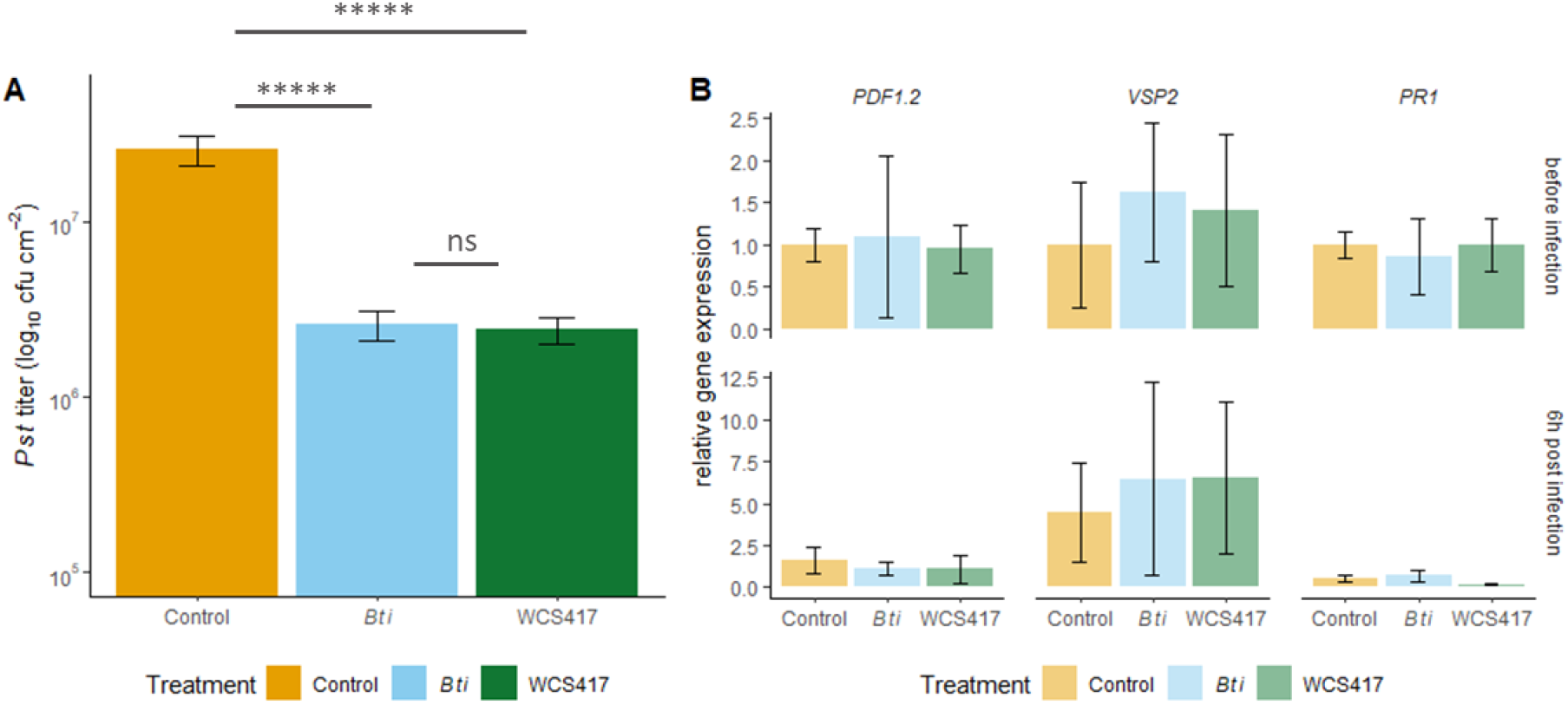
*Bacillus thuringiensis* var. *israelensis* (*Bti*) and *Pseudomonas simiae* WCS417r (WCS417) trigger induced systemic resistance (ISR) in *Arabidopsis thaliana* in the absence of SA and JA marker gene expression or priming. The roots of 10-day-old, sterile-grown *A. thaliana* seedlings were inoculated with *Bti* (blue bars), WCS417 (green bars), or a corresponding control solution (yellow bars). Following 3.5 weeks on soil, the leaves of the treated plants were infiltrated with *P. syringae* pathovar *tomato* (*Pst*). **(A)** *In planta Pst* titers at 4dpi. Bars represent the mean of three biologically independent experiments, including three replicates each ± SD. Asterisks indicate significant differences between the treatments indicated by the corresponding lines (pairwise Wilcoxon test, adjusted for multiple testing by Benjamini-Hochberg procedure, ****, *p* <0.0001; ns, not significantly different). **(B)** *PDF1*.*2, VSP2*, and *PR1* transcript accumulation in leaves of plants treated as in (A) and sampled before infection (upper panel) or 6 hours (h) after inoculation of the leaves with *Pst* (lower panel). Transcript accumulation was determined relative to that of *UBIQUITIN* by RT-qPCR. Bars represent mean values of three biologically independent experiments ± SD. Statistically significant differences were excluded using pairwise Wilcoxon test, adjusted for multiple testing by Benjamini-Hochberg procedure.

In contrast to SAR, which is classically associated with SA signalling, WCS417-induced ISR has previously been associated with JA signalling (Pieterse et al., 1996; Pieterse et al., 1998; Pozo et al., 2008). *PLANT DEFENSIN 1*.*2* (*PDF1*.*2*) and *VEGETATIVE STORAGE PROTEIN 2* (*VSP2*) are marker genes of the MYC2-independent and MYC2-dependent JA signalling pathways, respectively (Pieterse, Van der Does, Zamioudis, Leon-Reyes, & Van Wees, 2012). Here, we tested whether ISR induction leads to changes in JA signalling by conducting RT-qPCR analysis of *PDF1*.*2* and *VSP2* transcript accumulation. Additionally, we tested a possible influence of ISR on SA signalling targeting the SA marker gene *PATHOGENESIS RELATED 1* (*PR1*) (van Loon, Rep, & Pieterse, 2006). We sampled leaves of *Bti-*, WCS417-, and control-treated plants prior to a pathogenic challenge with *Pst* and 6 h post infection. *PDF1*.*2, VSP2*, and *PR1* transcript accumulation in *Bti-* as well as WCS417-treated plants was not significantly different in comparison to that in control-treated plants (Fig. 1B, upper panel). Thus, the induction of ISR did not induce transcript accumulation of these genes. Similarly, *PDF1*.*2, VSP2*, and *PR1* transcript accumulation was not different in ISR-induced as compared to control-treated plants sampled 6 h after challenge inoculation of the leaves with *Pst* (Fig. 1B, lower panel), indicating that transcript accumulation was not primed by either of the ISR treatments. Thus, under the experimental conditions used here, both *Bti* and WCS417 triggered ISR against *Pst* in *A. thaliana* in the absence of detectable induction or priming of JA and SA marker genes.

### Varying molecular requirements for WCS417- and *Bti-*induced ISR

WCS417-induced ISR has been shown to depend on functional MYC2-associated JA defences, but not on the accumulation of SA (Pieterse et al., 1996; Pieterse et al., 1998; Pozo et al., 2008). Here, we compared the functionality of ISR induced by *Bti* as compared to WCS417 in *A. thaliana* mutants with compromised JA defences *(jin1/myc2*) and also in mutants with compromised SA accumulation (*sid2*) and signalling (*npr1*). ISR was induced as described above, and the leaves of the plants were inoculated with *Pst*. Col-0 wild type supported less *Pst* growth in the leaves of plants pre-treated with either WCS417 or *Bti* as compared to the controls, confirming that ISR was induced in response to both bacterial strains (Fig. 2A). As reported before (Nickstadt, 2004), the *jin1* (*myc2*) mutant supported less *Pst* growth than Col- 0 wild type plants (Fig. 2A). In accordance with previous findings (Pozo et al., 2008), WCS417- induced ISR was compromised in *jin1* mutant plants (Fig. 2A). Similarly, *Bti-*induced ISR was abolished in *jin1* plants resulting in similar or slightly elevated growth of the *Pst* challenge inoculum as compared to that observed in control-treated plants. Thus, the *JIN1-*encoded, JA-associated transcription factor MYC2 is essential for ISR triggered by both WCS417 and *Bti*. Similarly, neither WCS417 nor *Bti* treatments reduced the *Pst* titers in the leaves of *sid2* or *npr1* mutant plants (Fig. 2A). This suggests that ISR induced by both WCS417 and *Bti* under the experimental conditions used here, depends on functional pathogen-induced SA accumulation and signalling.

**Figure 2.**
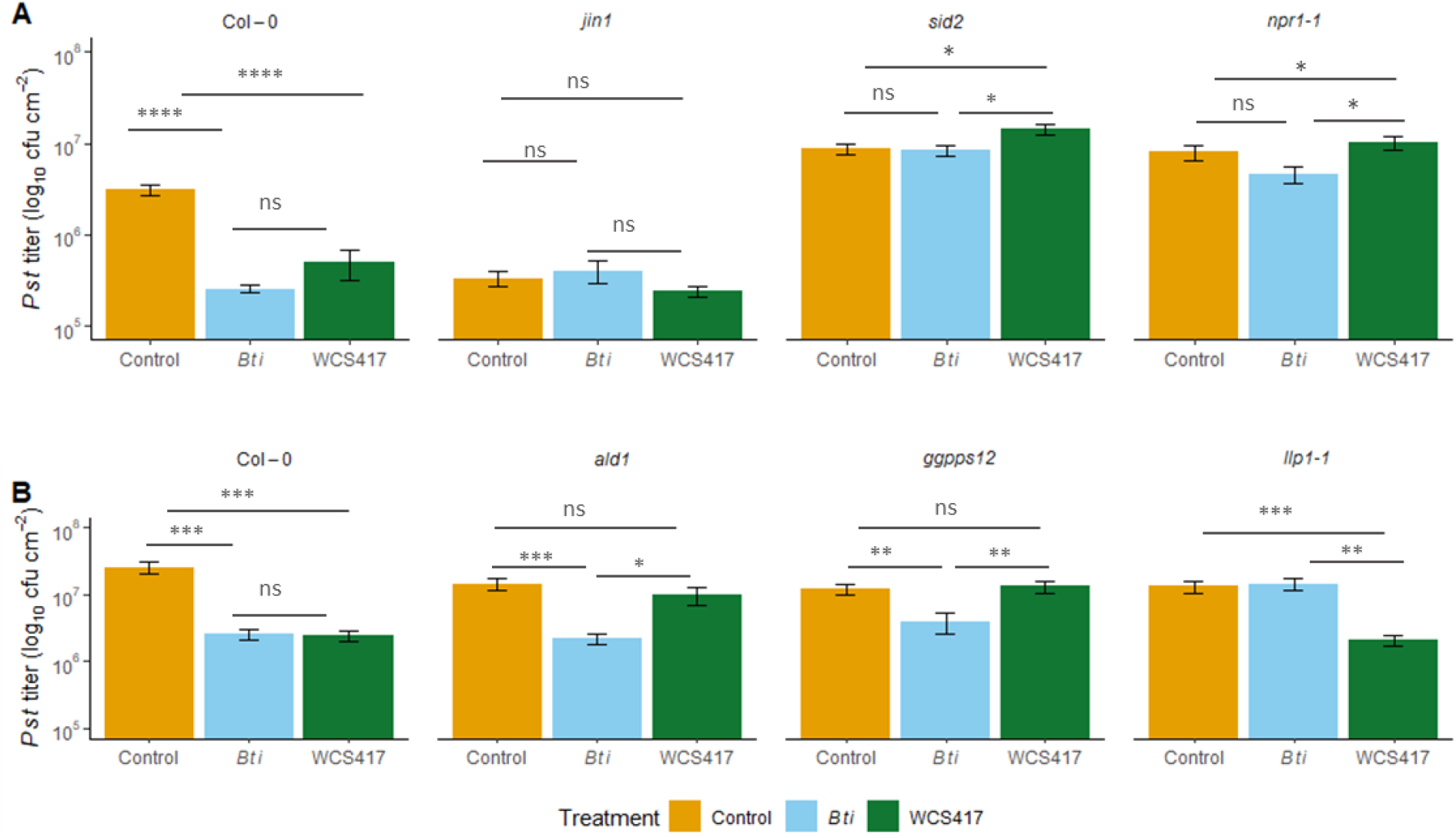
Characterization of the molecular requirements of *Bti-* and WCS417-triggered ISR. The roots of 10-day-old seedlings of the genotypes indicated above the panels were inoculated with *Bti* (blue bars), WCS417 (green bars), or a corresponding control solution (yellow bars). Following 3.5 weeks on soil, the leaves of the plants were inoculated with *Pst*. The resulting *in planta Pst* titers at 4 dpi are shown. Bars represent the mean of three biologically independent experiments, including three replicates each ± SD. Asterisks indicate significant differences between the treatments indicated by the corresponding lines (pairwise Wilcoxon test, adjusted for multiple testing by Benjamini-Hochberg procedure, *, *p* <0.05, **, *p* <0.01, ***, *p* <0.001, ****, *p* <0.0001; ns, not significantly different).

Recent evidence suggests roles of SAR-associated signalling intermediates in ISR (Cecchini, Steffes, Schlappi, Gifford, & Greenberg, 2015; Shine et al., 2019). Here, we assessed the involvement of Pip-dependent pathways in ISR by using *ald1* mutant plants with defects in Pip biosynthesis (Navarova et al., 2012). We also tested the involvement of SAR-associated volatile monoterpenes as well as their perception by monitoring ISR in the respective loss-of-function mutants *ggpps12* and *llp1* (Wenig et al., 2019). In comparison to the respective control treatments, treatment of *ald1* and *ggpps12* plants with *Bti* resulted in decreased *Pst* titers, suggesting that ISR had been induced in these plants (Fig. 2B). In contrast, treatment of the same mutants with WCS417 did not reduce growth of the *Pst* challenge inoculum, indicating that WCS417-triggered ISR was compromised in *ald1* and *ggpps12* mutant plants. This implies the involvement of Pip as well as monoterpenes in the realisation of immunity in WCS417- dependent ISR. In contrast to WCS417, *Bti* triggered ISR by a mechanism relying on the SAR-associated signalling factor LLP1: the *Pst* challenge inoculum grew to similar titers in the leaves of *Bti-*treated compared to control-treated *llp1* mutant plants (Fig. 2B). Taken together, the data suggest that WCS417 and *Bti* trigger ISR via two at least partially distinct mechanisms.

### Microbial composition of the phyllosphere differs in dependency of the ISR-eliciting bacterial strain

To address the question, whether the composition of the microbiome changes in leaves of plants undergoing ISR, we performed amplicon sequencing of the bacterial 16S rRNA gene. To this end, we treated plants at the age of 10 days with either *Bti*, WCS417, or MgCl_2_ as the control. 3½ weeks later, we harvested the leaves, isolated the DNA, and amplified and sequenced the regions V5-V7 of the 16S rRNA gene. In total 830.276 reads were sequenced. After data pre-processing (see Methods), the remaining 622.234 reads, averaging 36.602 reads per sample, were assigned to 1844 amplicon sequence variants (ASVs) acting as a proxy for bacterial species. Per sample 110 to 432 ASVs were identified. To assess the bias introduced by DNA isolation and PCR as well as to detect contaminations, we additionally processed a commercially available bacterial standard (ZYMO, see Methods). The microbial standard revealed no gross bias with respect to sequencing reads per bacterial group and only a slight contamination by a *Ralstonia* sp. (Supplementary Fig. S1). This data suggests that our samples were not subject to significant bias or contamination during DNA isolation as well as replication.

To obtain a general overview of the microbial composition of our samples, we first examined the sequencing data on the phylum level. Among the ten most abundant phyla were Proteobacteria, Firmicutes, Bacteroidetes and Actinobacteria (Fig. 3), which correspond to the phyla which were previously described as “core-phyla” for the microbiome of plants’ phyllosphere (Lundberg et al., 2012; Vorholt, 2012). Additionally, high counts of Cyanobacteria were detected, mainly caused by one ASV. This is rather unusual and presumably due to high air humidity during plant growth. The remaining phyla of high abundance where Myxococcota, Gemmatimonadota, Bdellovibrionota, Acidobacteria and Abditibacteriota. We detected differences in the phylum composition between the treatments. WCS417-treated plants contained more Bacteroidota and less Actinobacteria than control- and *Bti*-treated plants (Fig. 3).

**Figure 3.**
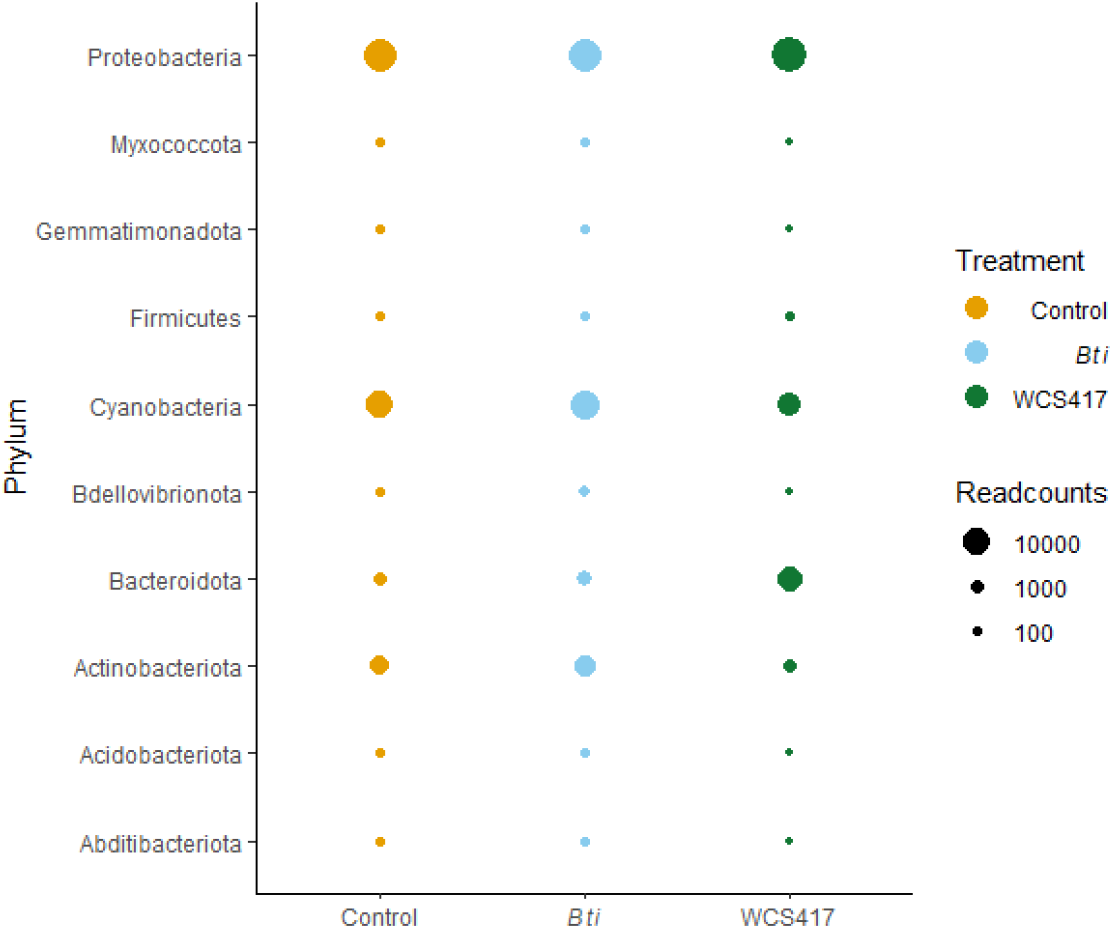
Distribution of 16S rRNA gene amplicon reads among the ten most abundant phyla in the *A. thaliana* phyllosphere microbiome of plants undergoing control (in yellow) and ISR-inducing treatments with *Bti* (in blue) or WCS417 (in green). Circle sizes represent mean read counts from five (*Bti* and WCS417) to six (control) biologically independent replicate experiments.

### The microbiome of WCS417-treated plants displays reduced species diversity

In the next step, we examined the ASVs by plotting a rarefaction curve of the amplicon data. The rarefaction curve confirmed a sufficient sequencing depth by showing a clear saturation of the curves (Fig. 4A). Additionally, the rarefaction curves revealed a significantly lower number of ASVs in WCS417-treated plants in comparison to *Bti-* or control-treated plants. The microbiomes of WCS417-treated plants on average contained 165 ASVs per sample in comparison to 331 or 361 ASV per sample in *Bti-* and control-treated plants, respectively (Fig. 4A). Therefore, we analysed ASV richness and evenness utilizing the Shannon’s Index (Spellerberg & Fedor, 2003) and dominance of single ASVs using the Simpson’s Index (Simpson, 1949). The apparent lower species richness in WCS417-treated plants was confirmed by the Shannon’s Index, which was significantly lower (p<0.05) in WCS417-treated plants than in *Bti-* or control-treated plants (Fig. 4B). The Simpson’s Index, which does not account for species richness but rather for dominance of single species, did not reveal significant differences between the treatments (Fig. 4B). Thus, species diversity was reduced in the leaf microbiome of WCS417-treated plants, while dominance by species was not different from that in *Bti*- and control-treated plants.

**Figure 4.**
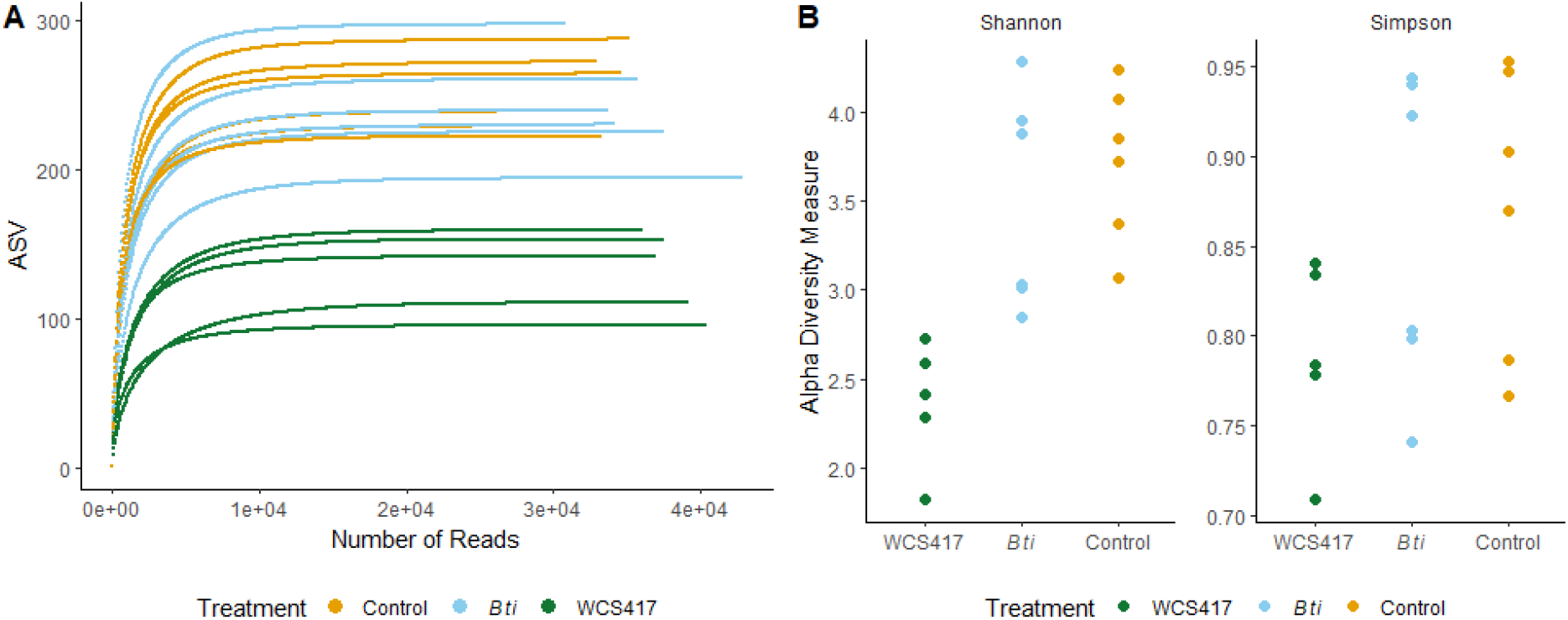
Diversity analysis of the phyllosphere microbiome of *A. thaliana* undergoing control (in yellow) and ISR-inducing treatments with *Bti* (in blue) or WCS417 (in green). **(A)** Rarefaction curves of the sequenced samples correlating the number of detected amplicon sequence variants (ASVs) on the Y-axis to the number of sequenced reads on the X-axis. Samples from WCS417-treated plants contain significantly fewer ASVs (pairwise Wilcoxon test, adjusted for multiple testing by Benjamini-Hochberg procedure, WCS417 – control, *p* = 0.013). **(B)** Shannon’s Index (left) and Simpson’s Index (right). The Y-axis represents the respective index value, and dots indicate the values of individual samples. Samples from WCS417-treated plants have a lower Shannon’s Index than *Bti*- or control-treated plants (pairwise Wilcoxon test, adjusted for multiple testing by Benjamini-Hochberg procedure, WCS417 – control, Shannon: *p* =0.013, Simpson: *p*= 0.38).

### Nonmetric multidimensional scaling reveals differences and similarities of bacterial composition between the different treatments

In the next step, we addressed the question, how similar or distinct the different samples are with regards to their composition under consideration of the relative relatedness of the different ASVs. To this end, we calculated weighted Unifrac-distances (Lozupone & Knight, 2005) and performed a nonmetric multidimensional scaling to see whether the samples cluster for example by treatment or replicate number. The non-metric multidimensional scaling shows that the microbiome of plants treated with WCS417 clustered distinctly from that of *Bti*- and control-treated plants (Supplementary Fig. S2). Despite the similar clustering between the *Bti* and control treatments, both groups of samples clustered in a significantly different manner. In some samples we observed a slight clustering according to the experimental replicate (e.g., number 6 and number 7). This hints at possible batch effects due to treatment, growth, or sampling of the plants.

### Occurrence of ISR-eliciting bacterial strains on the ISR-treated plants

We wanted to analyze if the bacterial strains we used to elicit ISR also occurred on the leaves of the plants. Therefore, we examined the absolute numbers of ASV3 and ASV953, whose 16S rRNA gene sequences correspond to that of WCS417 and *Bti*, respectively. In WCS417- treated plants, the reads of WCS417 on the leaves make up ∼25% of the reads per sample, averaging nearly 10.000 reads per sample (Fig. 5A). This suggests a possible contamination of the phyllosphere with WCS417. Alternatively, it is possible that WCS417 actively proliferates on *A. thaliana* leaves. In support of the latter hypothesis, we detected moderate growth of a WCS417 inoculum in *A. thaliana* leaves (Supplementary Fig. S3). While absent from control-treated plants, WCS417 was found with up to 100-1000 reads per sample in leaves of *Bti*- treated plants (Figure 5A), suggesting a possible recruitment of WCS417 to the phyllosphere of *Bti-*treated plants. In *Bti-*treated plants, *Bti* was detected with 7 reads in a single sample (Fig. 5A). Also, 14 reads corresponding to *Bti* were detected in one sample from control-treated plants. This finding suggests that contamination of the phyllosphere during the ISR-inducing treatment had been negligible. Upon inoculation of *A. thaliana* leaves with *Bti*, we observed that the titers stagnated over time (Supplementary Fig. S3), suggesting that *Bti* does not proliferate in *A. thaliana* leaves. Taken together, inoculation of *A. thaliana* roots with ISR-inducing bacteria resulted in proliferation of WCS417, but not *Bti*, on the leaves of the treated plants. Strikingly, the data suggest that *Bti-*triggered ISR was accompanied by the recruitment of WCS417 to the phyllosphere.

**Figure 5.**
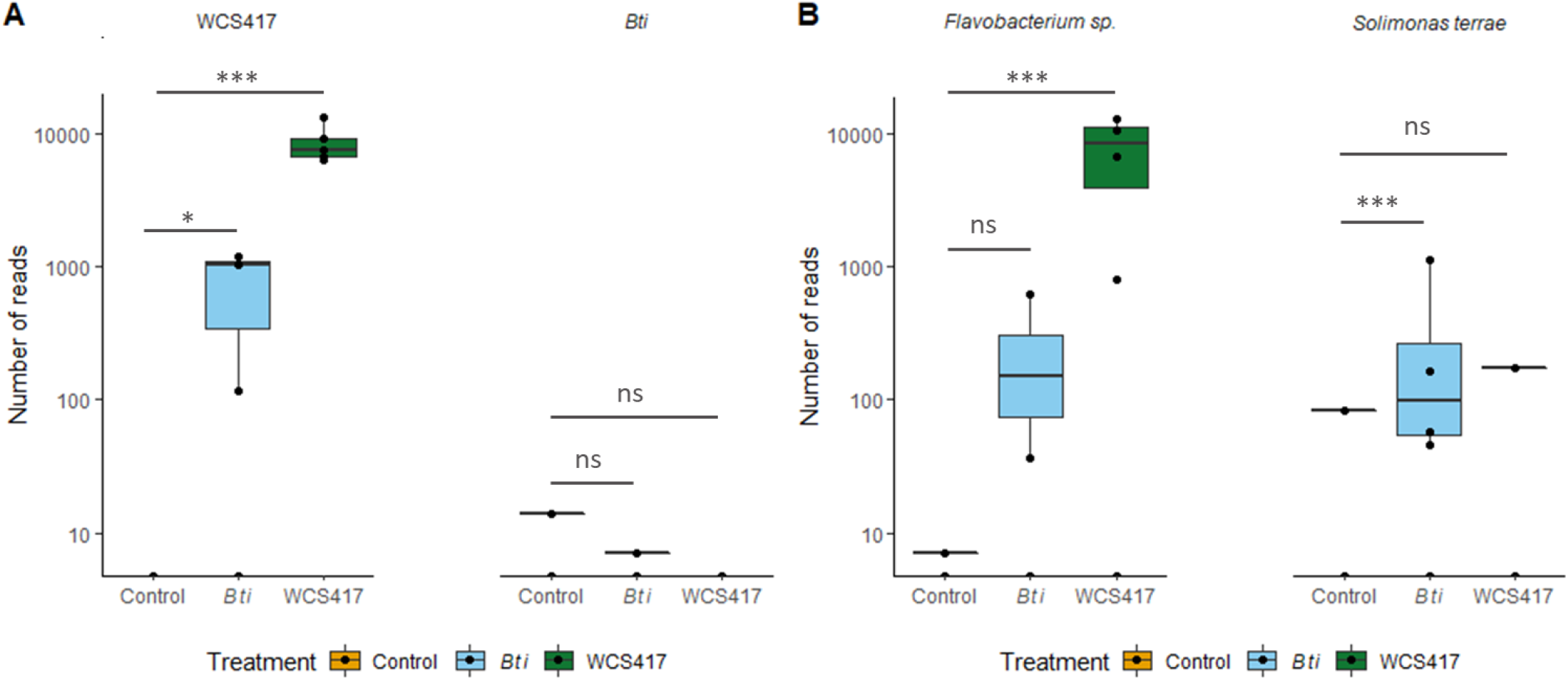
Abundance of distinct bacterial species in the phyllosphere microbiome of *A. thaliana* undergoing control (in yellow) and ISR-inducing treatments with *Bti* (in blue) or WCS417 (in green). Boxplots indicate average numbers of sequenced reads corresponding to the species indicated above the panels from five (*Bti* and WCS417) to six (control) samples after normalization to total read counts per sample. Asterisks indicate significant differences between the treatments indicated by the corresponding lines (pairwise Wilcoxon test, adjusted for multiple testing by Benjamini-Hochberg procedure, *, *p* <0.05, **, *p* <0.01, ***, *p* < 0.001).

### Different bacterial groups are enriched in the phyllosphere depending on the treatment

For the analysis of ASVs that appeared in significantly different abundance between treatments, only ASVs that were detected in at least three samples per treatment were taken into account. We utilized the R package “dada2” which was originally created for the analysis of RNAseq-data. This library has the advantage of utilizing more suitable methods for data normalization than the package “Phyloseq”, which we used for most of the remaining data analysis. In this manner, data normalization was independent of subsampling and the associated loss of data (McMurdie & Holmes, 2014).

In comparison to control-treated plants, *Bti-*treated plants displayed differential abundance of 14 ASVs and WCS417-treated plants of 42 ASVs (Supplementary Table S2). Most of the differentially accumulating ASVs were less abundant in ISR-treated compared to control-treated plants. Also, most of the significantly different ASVs were detected at relatively low read count numbers of 1000 reads in total or less. In contrast, two bacterial species were considerably enriched in the ISR-treated plants. In samples from WCS417-treated plants, a *Flavobacterium sp*. was detected at an average of 6000 reads per sample, while the same strain was detected with an average of 110 reads per sample in *Bti*-treated plants (Fig. 5B). By comparison, the same ASV was detected with 1 read in 1 control sample, and thus remained negligible on control-treated plants (Fig. 5B). Similarly, *Bti*-treated plants displayed a significant accumulation of a *Solimonas terrae* strain, which was detected in 4 out of 5 samples with an average of 231 reads per sample (Fig. 5B). The same ASV occurred in 1 sample each from control- and WCS417-treated plants. Taken together, ISR triggered by WCS417 and *Bti* was associated with enrichment of the phyllosphere microbiome with WCS417 and *Flavobacterium sp*., while *Bti* treatment additionally resulted in the enhanced recruitment of *S. terrae* to the *A. thaliana* phyllosphere.

### Microbe-microbe-host interactions in the *A. thaliana* phyllosphere

Because WCS417 proliferated in *A. thaliana* leaves, we tested if this proliferation triggered systemic immunity against *Pst*. To this end, we infiltrated the first true leaves of 4-5-week-old *A. thaliana* plants with WCS417 or with MgCl_2_ as the negative control. As a positive control, we used the bacterial strain *Pst/AvrRpm1* which is known to cause SAR (Breitenbach et al., 2014). Three days later, we performed a challenge infection of the systemic leaves with *Pst*. In comparison to the negative control treatment, *Pst/AvrRpm1* pre-treatment significantly decreased *Pst* propagation, indicating that SAR had been induced (Supplementary Fig. S4). In contrast, a local WCS417 leaf inoculation did not decrease *Pst* titers in the systemic tissue of the treated plants. Similar experiments using *Bti* as the primary, SAR-inducing treatment showed that a local leaf inoculation with *Bti* triggered SAR in systemic leaves (Supplementary Fig. S4). Thus, when inoculated onto *A. thaliana* leaves, *Bti*, but not WCS417, induced systemic immunity.

We next investigated if WCS417 locally enhances the immunity of *A. thaliana* leaves against *Pst*. Because the relative abundance of *Flavobacterium sp*. was significantly enhanced on the leaves of ISR-induced plants, we also tested if this bacterium affects defence. To this end, we used bacterial strain Leaf82 from the *At*-LSPHERE collection (Bai et al., 2015), which displays 100% sequence identity of its V5-V7 16S rRNA gene region with that of *Flavobacterium sp*. To study induced resistance, WCS417 and Leaf82 were syringe-infiltrated into leaves of 4-5- week-old *A. thaliana* plants. Two days later, the same leaves were infiltrated with *Pst*. In comparison to the control treatment, WCS417 treatment of the leaves caused a reduction of *Pst* proliferation (Fig. 6A), suggesting that WCS417 locally induced resistance on *A. thaliana* leaves. Because WCS417-induced resistance was not observed in *npr1* mutant plants (Fig. 6A), the observed reduction of *Pst* growth was likely associated with plant immunity. In contrast, Leaf82 treatment did not reduce *Pst* proliferation on the leaves and thus did not enhance plant defences against this pathogen (Fig. 6B).

**Figure 6.**
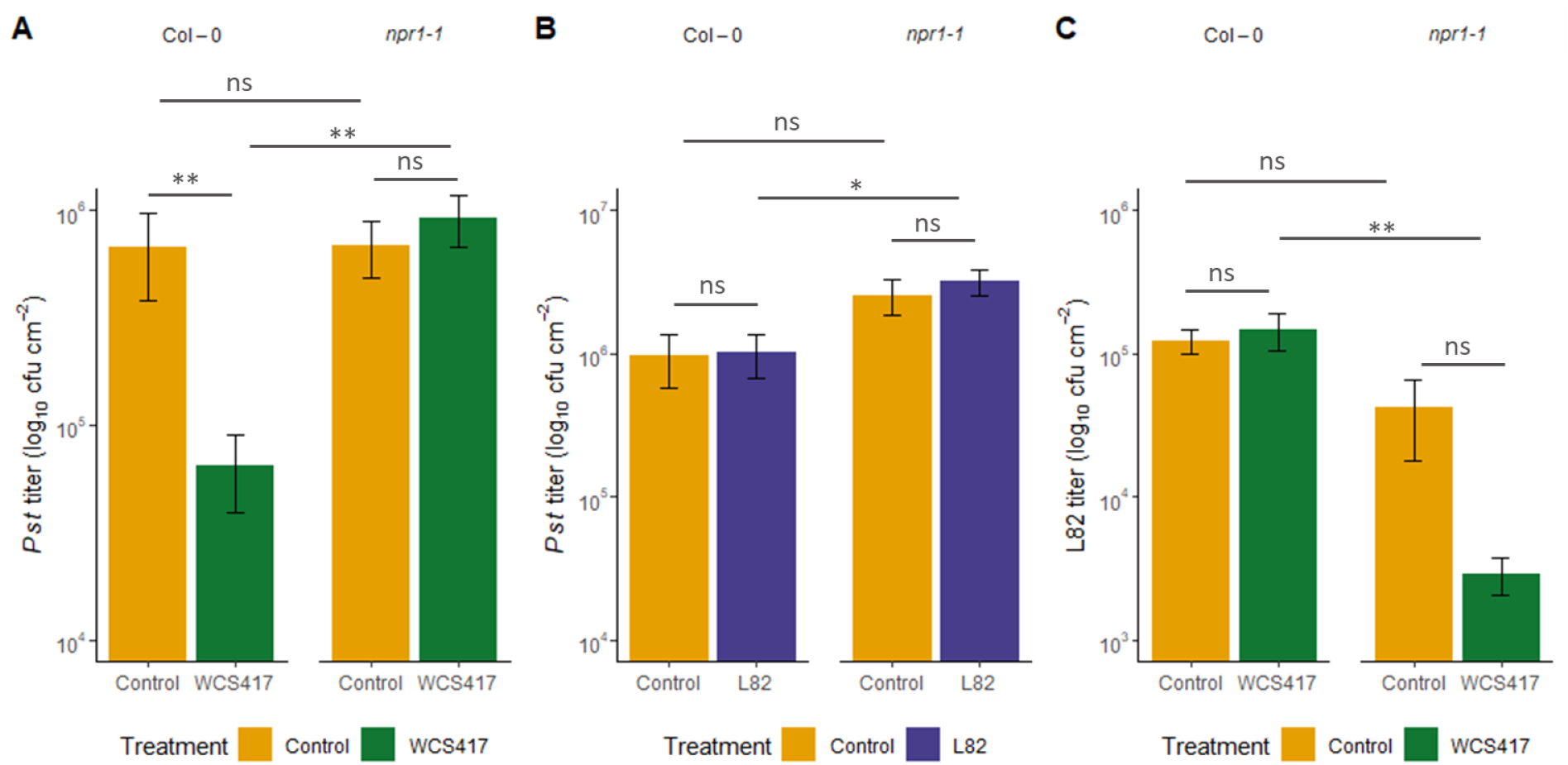
Local plant-microbe-microbe interactions. Leaves of 4-5-week-old Col-0 wild type and *npr1* mutant plants (as indicated above the panels) were infiltrated with WCS417 (green bars in A/C), *At-*L-Sphere *Flavobakterium sp*. Leaf82 (L82; purple bars in B), or a corresponding control solution (yellow bars in A-C). Two days later, the same leaves were infiltrated with *Pst* (A/B) or Leaf82 (C), titers of which were determined at 4 dpi. Bars represent average *in planta Pst* (A/B) and Leaf82 (C) titers from 6 to 9 samples derived from two (C) to three (A/B) biologically independent experiments ± SD. Asterisks indicate significant differences between the treatments indicated by the corresponding lines (pairwise Wilcoxon test, adjusted for multiple testing by Benjamini-Hochberg procedure, *, *p* <0.05, **, *p* <0.01; ns, not significantly different).

Finally, we studied if the local WCS417-induced defence response of *A. thaliana* influenced the proliferation of Leaf82. To this end, we performed the same induced resistance experiment as above. Two days after treating leaves with WCS417 or a control solution, we infiltrated the same leaves with Leaf82. Although WCS417-induced ISR was associated with enhanced proliferation of *Flavobacterium sp*. on the leaves, leaf inoculation of WCS417 did not cause enhanced growth of a subsequent Leaf82 inoculum (Fig. 6C). Although it thus seems as though WCS417 does not affect Leaf82 through plant responses, the proliferation of Leaf82 was reduced on *npr1* mutant as compared to wild type plants (Fig. 6C). Similarly, WCS417 proliferated less on *npr1* mutant than on wild type plants (Supplementary Fig. S5). Taken together, the data suggest that WCS417 activates *NPR1-*dependent responses in plants that reduce growth of pathogenic *Pst* and at the same time enhance WCS417 proliferation. Because *Flavobacterium sp*. or Leaf82 titers on leaves appear to correlate with those of WCS417 in various treatments (Figs. 5, 6C, and S5), the data suggest that this bacterium is under direct influence of WCS417 and thus, during ISR, subject to plant-microbe-microbe interactions.

## Discussion

We showed here that *Bacillus thuringiensis israelensis* (*Bti*) triggers ISR in *A. thaliana*. Additionally, local leaf application of *Bti* enhanced immunity in systemic leaves of the treated plants. Until now, *Bti* has been known mainly for its CRY-proteins, which are toxic specifically for insects and are widely used as crop protection agents in agriculture (Bravo, Gill, & Soberón, 2007). ISR-eliciting properties of *Bacillus thuringiensis* subspecies have so far been observed in tomato (Hyakumachi et al., 2013; Raddadi et al., 2007; Takahashi et al., 2014). Our data suggest that *Bti* enhances systemic immunity in *A. thaliana* by inducing both root-to-leaf and leaf-to-leaf systemic immune signalling.

*Bti-*triggered ISR depends on functional JA and SA signalling and on LLP1, which is also essential for SAR (Breitenbach et al., 2014). ISR triggered by the model strain WCS417 depended on SA and JA signalling and further on Pip and monoterpene biosynthesis (Figure 2). Until recently, SAR and ISR were believed to depend on different molecular mechanisms. Recent evidence, however, suggests that a number of SAR-associated defence cues also promotes ISR (Vlot et al., 2020). These cues include azelaic acid, AZELAIC ACID INDUCED1, and glycerol-3-phosphate (Cecchini et al., 2019; Cecchini et al., 2015; Shine et al., 2019), which act downstream of Pip in SAR (Wang et al., 2018). Here, Pip promoted ISR triggered by WCS417, but not *Bti*. Pip and glycerol-3-phosphate further cooperate to drive monoterpene emissions during SAR (Wenig et al., 2019). Consequently, monoterpene emissions promoted the same WCS417-triggered ISR mechanism as Pip. Although LLP1 promotes SAR in a positive feedback loop with Pip (Wenig et al., 2019), this function of LLP1 does not appear involved in WCS417-triggered ISR. It is possible that another function of LLP1, which we previously connected with local release of SAR-associated long-distance signals (Wenig et al., 2019), is important for ISR triggered by *Bti*. Together, this work suggests functions of three SAR-associated signalling intermediates in ISR, supporting the hypothesis that the mechanisms of SAR and ISR are not as different as traditionally believed.

Under our experimental conditions, WCS417 appears to be recruited to the phyllosphere, when applied to roots (Figure 5). There, the bacteria proliferate (Supplementary Fig. S3), and potentially enhance the immunity of the leaves against *Pst* via a local induced resistance response (Figure 6). This contrasts with findings of Pieterse et al. (1996, 1998) who showed that WCS417 remains confined to the rhizosphere during the elicitation of ISR. WCS417- triggered ISR further was functional in SA-deficient *NahG* plants (Pieterse et al., 1996), whereas the same response was compromised in *sid2* plants with reduced pathogen-induced SA biosynthesis (Figure 2)(Wildermuth et al., 2001). These contrasting results could be a consequence of the differential WCS417 proliferation on leaves in our studies. Alternatively, the combined data suggest a possible role of low remaining SA levels or SA-derivatives such as MeSA, which accumulate in NahG plants, in WCS417-triggered ISR (Lim et al., 2020; Park, Kaimoyo, Kumar, Mosher, & Klessig, 2007). In accordance with previous findings (Pozo et al., 2008), the WCS417-triggered ISR response further relied on functional MYC2-dependent JA signalling (Figure 2), suggesting synergism between SA and JA in ISR-activated leaves.

ISR not only affects the plant itself, but also seems to change the habitat it provides in its phyllosphere, leading to changes in the microbial composition of the leaf. Here, ISR was associated with a higher relative abundance of WCS417 and *Flavobacterium sp*. in the *A. thaliana* phyllosphere. *Bti* additionally triggered a distinct enrichment of a *Solimonas terrae* strain first isolated in soil from Korea (S.-J. Kim et al., 2014). Until now, not much is known about this and the other five known species of the genus *Solimonas*. Recently, *Solimonas terrae* was associated with changes in the microbiome of plants after a growth-stimulating cold plasma treatment in *A. thaliana* (Tamošiūnė et al., 2020). Here, proliferation of *S. terrae* appeared uniquely related to the ISR trigger *Bti*, and thus might be responsive to true systemic signalling. In future, it will be of interest to investigate possible beneficial properties of this bacterial strain for plant health.

In the phyllosphere of WCS417-treated plants, WCS417 and *Flavobacterium sp*. together accounted for more than 50% of the sequenced reads. Consequently, the relative abundance of other bacterial strains was reduced, possibly because they were supplanted by WCS417 and *Flavobacterium sp*. Species belonging to the phylum Proteobacteria have been proposed to act as ‘key-stone’ bacterial species in phyllosphere microbiomes (Carlstrom et al., 2019). Single ‘key-stone’ strains can have significant effects on the overall microbial composition. However, Leaf82, the strain highly similar to the *Flavobacterium sp*. we found enriched in association with WCS417, does not appear to be significantly influenced by Gammaproteobacteria related to WCS417 (Carlstrom et al., 2019). Here, consecutive leaf inoculations of WCS417 and Leaf82 suggest that Leaf82 proliferation is influenced by WCS417, suggesting direct microbe-microbe interactions between these strains in the *A. thaliana* phyllosphere.

Notably, the phyllosphere microbiome of WCS417-treated plants displayed a significantly reduced species richness. Because lower richness in microbiota has been associated with a lower stability of the microbiome and a higher risk of dominance by pathogens (Chaudhry et al., 2020; Tao Chen et al., 2020; Ives & Hughes, 2002), this suggests a possible trade-off of ISR in plants. In this respect, it seems of interest that we detected considerably less significant shifts in the phyllosphere microbiome composition than Chen et al. (2020), who studied the influence of local immune response driven by pathogen-associated molecular patterns (PAMPs). The comparatively moderate phyllosphere microbiome changes in response to ISR likely reflect the fact that ISR is established as a form of priming (U. Conrath, G. J. Beckers, C. J. Langenbach, & M. R. Jaskiewicz, 2015; Mauch-Mani et al., 2017). During priming, the bulk of defence-associated molecular responses do not become evident before a pathogen challenge (Martinez-Medina et al., 2016). Therefore, it is not unexpected that the microbiome also displays only a moderate response to the induction of ISR. By comparison, *A. thaliana* mutants, which were defective in the MIN7-vesicle-trafficking pathway and incapable of mounting PAMP-triggered immunity (PTI), displayed more significant shifts in the relative abundance of Proteobacteria (up) and Actinobacteria (down). Deployment of this “incorrectly” assembled microbiome onto gnotobiotic plants led to necrosis and stunted plant growth (T. Chen et al., 2020). These findings associate compromised PTI responses with reduced plant fitness, caused by changes in the phyllosphere microbiome. In our experiments, although WCS417 reduced the species richness of the phyllosphere microbiome, defence against *Pst* was enhanced. Notably, we focused on the bacterial part of the microbiome, and cannot exclude possible additional effects of e.g. fungi and other eukaryotic microbes (Chaudhry et al., 2020).

Recruitment of WCS417 to the phyllosphere appears to be an active process driven by NPR1-mediated plant immunity. Data shown in Figure 6 suggest that enhanced proliferation of WCS417, in turn, drives proliferation of *At*-LSPHERE *Flavobacterium sp*. Leaf82. Species belonging to the genus *Flavobacterium* are known to be well-adapted to the phyllosphere, living epi-as well as endophytically on *A. thaliana* plants (Bodenhausen, Horton, & Bergelson, 2013). They are capable of metabolizing complex carbon-sources such as pectin, hemicellulose, and peptidoglycan components of gram-positive cell-walls of bacteria (Kolton, Sela, Elad, & Cytryn, 2013; Peterson, Dunn, Klimowicz, & Handelsman, 2006). By those attributes, *Flavobacteria* spp. can outcompete other bacterial groups. Additionally, they have been assigned enhanced biocontrol as well as plant growth-promoting properties. *Flavobacterium spp*., for example, produce cyanide acting as an antimicrobial agent as well as compounds that act as plant growth-promoting hormones, including auxins, gibberellins and cytokinins (Gunasinghe, Ikiriwatte, & Karunaratne, 2004; Hebbar, Berge, Heulin, & Singh, 1991; Maimaiti et al., 2007; Sang & Kim, 2012). It is thus conceivable that ISR triggers the recruitment of plant growth-promoting microbiota to the phyllosphere.

## Conclusions

ISR triggered in *A. thaliana* by *Bti* or WCS417 leads to the recruitment of microbiota with plant growth-promoting properties to the phyllosphere. This recruitment depends on interconnected plant-microbe and microbe-microbe interactions. WCS417-triggered ISR reduces the species richness of the phyllosphere microbiome, which hints at a possible trade-off of ISR in plants. Whereas short term effects did not appear to enhance plant disease susceptibility, further investigations are necessary to gain insights into the long-term effects of these plant-microbe-microbe interactions on plant health.

## Acknowledgements

The authors thank Dr. Michael Rothballer (Institute of Network Biology, Helmholtz Zentrum Muenchen, Germany) for providing *Bacillus thuringiensis* var. *israelensis* and Dr. Julia Vorholt (ETH Zürich, Switzerland) for providing *At-*LSPHERE Leaf82 and helpful advice on this work. This work was funded by the DFG as part of the priority program DECRyPT (to MS and ACV).

## Supplementary Materials

**Supplementary Table S1** Primers used for RT-qPCR

**Supplementary Table S2** Amplicon sequence variants (ASVs) with a significantly different relative abundance in the phyllosphere microbiome of *Bti-* or WCS417-treated plants as compared to control-treated plants.

**Supplementary Figure S1** 16S rRNA gene amplicon sequencing results of the microbial standard control

**Supplementary Figure S2** Microbiome composition analysis of the phyllosphere of *A. thaliana* undergoing control and ISR-inducing treatments

**Supplementary Figure S3** Growth of ISR-inducing bacteria in *A. thaliana* leaves.

**Supplementary Figure S4** Systemic immunity in response to local leaf application of *Bti* and WCS417

**Supplementary Figure S5** WCS417 titers in Col-0 (wild type) and *npr1-1* mutant plants.

## Notes

Funding: This work was funded by the DFG as part of priority program SPP 2125 (to MS and ACV).

### Competing Interest Statement

The authors have declared no competing interest.

