## Supplementary Material for "Induced systemic resistance impacts the phyllosphere microbiome through plant-microbe-microbe interactions"

**Supplementary Table S1.** Primers used for RT-qPCR

| Primer name | Sequence (5'→3') |
|---|---|
| Ubiquitin F | AGATCCAGGACAAGGAGGTATTC |
| Ubiquitin R | CGCAGGACCAAGTGAAGAGTAG |
| PDF1.2 F | CCAAGTGGGACATGGTCAG |
| PDF1.2 R | ACTTGTGTGCTGGGAAGACA |
| PR1 F | CTACGCAGAACAACTAAGAGGCAAC |
| PR1 R | TTGGCACATCCGAGTCTCACTG |
| VSP2 F | GTTAGGGACCGGAGCATCAA |
| VSP2 R | AACGGTCACTGAGTATGGGT |

**Supplementary Table S2**. Amplicon sequence variants (ASVs) with a significantly different relative abundance in the phyllosphere microbiome of *Bti*- or WCS417-treated plants as compared to control-treated plants. ns, not significant

| ASV | Genus | $log_2$ fold change *Bti* | *p*-value *Bti* | $log_2$ fold change WCS417 | *p*-value WCS417 | number of samples ASV is present in | sum of all reads |
|---|---|---|---|---|---|---|---|
| ASV3 | Pseudomonas | 8,7862 | <0.05 | 16,5712 | < 0.0005 | 8 | 54671 |
| ASV4 | Flavobacterium | 6,2659 | ns | 20,5589 | < 0.0005 | 7 | 38331 |
| ASV17 | Pseudomonas | -3,1333 | ns | -7,9596 | <0.005 | 13 | 7196 |
| ASV39 | Class: Cyanobacteria | -1,0111 | ns | 10,8304 | <0.05 | 5 | 1901 |
| ASV41 | Pseudonocardia | -5,4500 | ns | -11,9146 | <0.05 | 6 | 2258 |
| ASV45 | Solimonas | 15,6612 | < 0.0005 | 8,9117 | ns | 6 | 1943 |
| ASV57 | Chryseobacterium | 2,3738 | ns | 14,6927 | <0.005 | 3 | 1468 |
| ASV93 | Rhodopseudomonas | -4,0275 | ns | -8,3072 | <0.05 | 7 | 928 |
| ASV106 | Nocardioides | 9,7871 | ns | 12,5552 | <0.05 | 5 | 730 |
| ASV181 | Kribbella | -8,1963 | ns | -17,8013 | < 0.0005 | 4 | 242 |
| ASV225 | Jatrophihabitans | 4,0864 | ns | -16,4214 | < 0.0005 | 7 | 233 |
| ASV237 | Bosea | 19,4199 | < 0.0005 | 30,0000 | < 0.0005 | 3 | 233 |
| ASV239 | Pseudonocardia | 0,7281 | ns | -19,0794 | < 0.0005 | 7 | 221 |
| ASV270 | Order: Burkholderiales | 3,6172 | ns | -14,7184 | <0.005 | 5 | 76 |
| ASV326 | Allorhizobium-Neorhizobium-Pararhizobium-Rhizobium | 28,9100 | < 0.0005 | 21,5489 | < 0.0005 | 3 | 141 |
| ASV343 | Burkholderia-Caballeronia-Paraburkholderia | -3,0481 | ns | -19,5871 | < 0.0005 | 5 | 131 |
| ASV348 | Pedosphaera | 2,6113 | ns | -16,2309 | < 0.0005 | 6 | 128 |
| ASV356 | Luteimonas | 29,1442 | < 0.0005 | 29,9840 | < 0.0005 | 3 | 124 |
| ASV362 | Gemmatimonas | -36,1374 | < 0.0005 | -8,2986 | ns | 3 | 96 |
| ASV380 | Burkholderia-Caballeronia-Paraburkholderia | 8,6898 | ns | -19,0662 | < 0.0005 | 3 | 110 |
| ASV398 | Family: Polyangiaceae | -4,5516 | ns | -16,4222 | < 0.0005 | 6 | 85 |

| ASV | Genus | $log_2$ fold change *Bti* | *p*-value *Bti* | $log_2$ fold change WCS417 | *p*-value WCS417 | number of samples ASV is present in | sum of all reads |
|---|---|---|---|---|---|---|---|
| ASV416 | Familiy: Solirubrobacteraceae | -7,2479 | ns | -14,5038 | <0.05 | 3 | 45 |
| ASV423 | Dokdonella | -7,0014 | ns | -19,0904 | < 0.0005 | 3 | 49 |
| ASV435 | Aquabacterium | -21,4329 | < 0.0005 | 8,3044 | ns | 4 | 86 |
| ASV441 | Marmoricola | 27,2299 | < 0.0005 | 28,2533 | < 0.0005 | 3 | 90 |
| ASV462 | Micropepsis | 2,2356 | ns | -14,3879 | <0.05 | 5 | 84 |
| ASV467 | Order: Elsterales | -8,6829 | ns | -21,0741 | < 0.0005 | 4 | 64 |
| ASV481 | Order: Flavobacteriales | 1,4597 | ns | -16,9788 | <0.005 | 5 | 67 |
| ASV491 | Devosia | 4,6399 | ns | -14,3560 | <0.05 | 6 | 66 |
| ASV562 | Family: Rhodospirillaceae | -8,0415 | ns | -15,5264 | <0.005 | 3 | 60 |
| ASV575 | Class: Acidimicrobiia | -2,2550 | ns | -17,5413 | < 0.0005 | 5 | 57 |
| ASV579 | Bdellovibrio | -7,4353 | ns | -22,7484 | < 0.0005 | 5 | 56 |
| ASV604 | Gaiella | -7,1387 | ns | -16,7629 | <0.005 | 4 | 52 |
| ASV642 | Finegoldia | -21,1282 | < 0.0005 | 8,8718 | ns | 4 | 46 |
| ASV663 | Order: Sphingobacteriales | 1,3680 | ns | -14,3434 | <0.05 | 3 | 43 |
| ASV668 | Anaeromyxobacter | -7,1207 | ns | -16,0941 | <0.005 | 3 | 40 |
| ASV669 | Novosphingobium | -7,3750 | ns | -22,7162 | < 0.0005 | 3 | 43 |
| ASV690 | Jatrophihabitans | 7,6648 | ns | -16,2582 | <0.005 | 3 | 41 |
| ASV708 | Bauldia | -23,4135 | < 0.0005 | -6,5078 | ns | 3 | 39 |
| ASV722 | Family: Phycisphaeraceae | -6,7048 | ns | -15,2851 | <0.005 | 3 | 38 |
| ASV763 | Phenylobacterium | 24,8175 | < 0.0005 | 26,6086 | < 0.0005 | 3 | 33 |
| ASV803 | Class: Cyanobacteria | 29,5048 | < 0.0005 | 23,8963 | < 0.0005 | 3 | 30 |
| ASV826 | Puia | -6,7607 | ns | -24,0423 | < 0.0005 | 3 | 29 |
| ASV870 | Class: Polyangia | 5,9490 | ns | -13,3471 | <0.05 | 4 | 27 |
| ASV898 | Family: Paracaedibacteraceae | 22,7797 | < 0.0005 | 29,8082 | < 0.0005 | 4 | 25 |
| ASV902 | Arthrobacter | 16,2890 | <0.005 | 24,0136 | < 0.0005 | 3 | 25 |

**A**

Composition of the microbial standard:

- *Bacillus subtilis*
- *Enterococcus faecalis*
- *Escherichia coli*
- *Lactobacillus fermentum*
- *Listeria monocytogenes*
- *Pseudomonas aeruginosa*
- *Salmonella enterica*
- *Staphylococcus aureus*

**B**

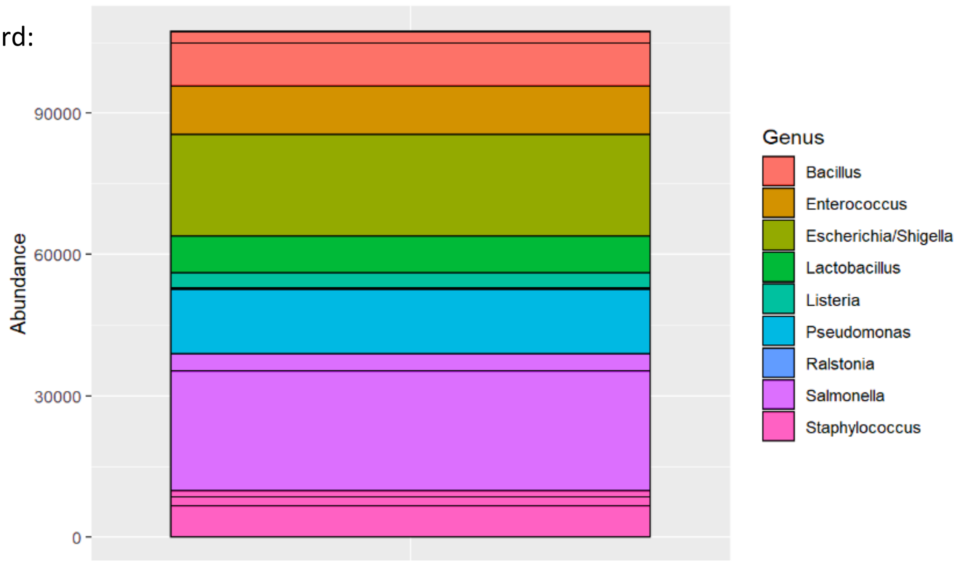


**Supplementary Figure S1**. 16S rRNA gene amplicon sequencing results of the microbial standard control. **(A)** The microbial standard (ZymoBIOMICS) used to assess sample contamination and sequencing bias contains the listed bacterial strains in equal amounts. **(B)** The standard was processed as an additional sample in the microbiome analysis (see methods). Y-axis represents absolute read counts, colours indicate genera as indicated on the right, and rectangles in the main diagram indicate different amplicon sequence (ASVs). We detected small amounts of one *Ralstonia sp.* (8 reads) as the only contaminant.

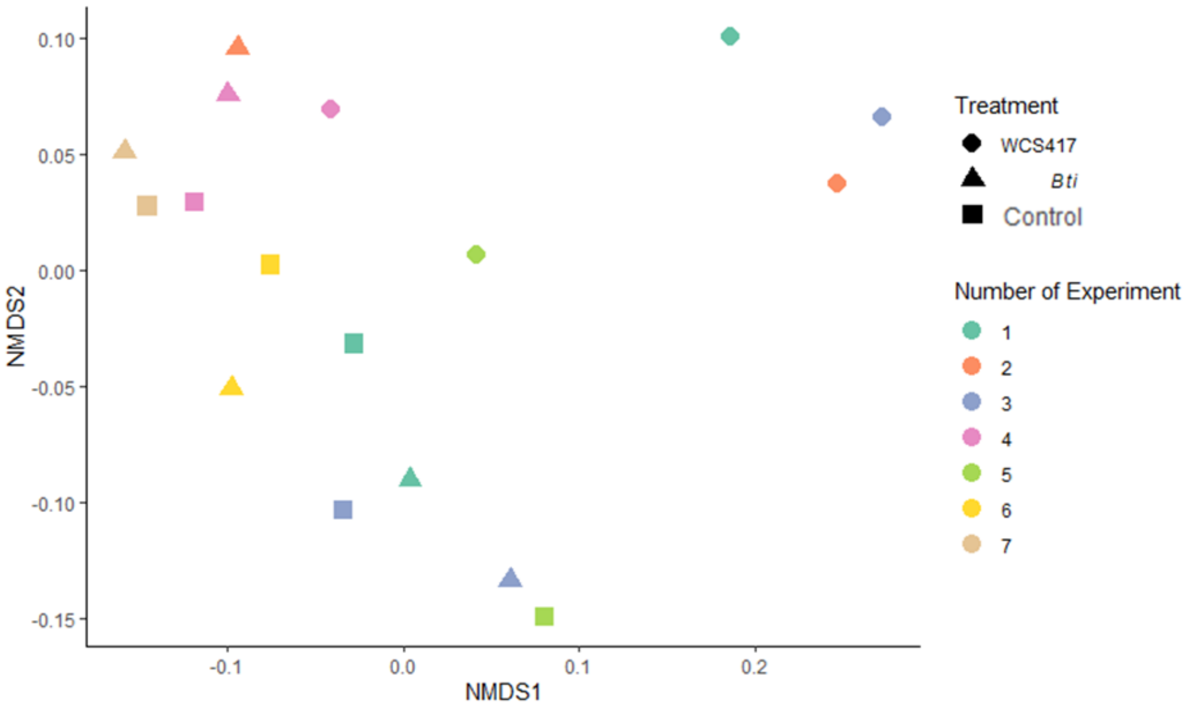


**Supplementary Figure S2**. Microbiome composition analysis of the phyllosphere of *A. thaliana* undergoing control (Control; squares) and ISR-inducing treatments with *Bti* (Bti; triangles) or WCS417 (417; circles). Different colours indicate different replicate experiments (numbers indicated on the right). Weighted Unifrac distances take the relative relatedness of microbes of the microbial community into account as well as the abundance of organisms to calculate a distance matrix. This matrix was visualized using nonmetric multidimensional scaling (NMDS).

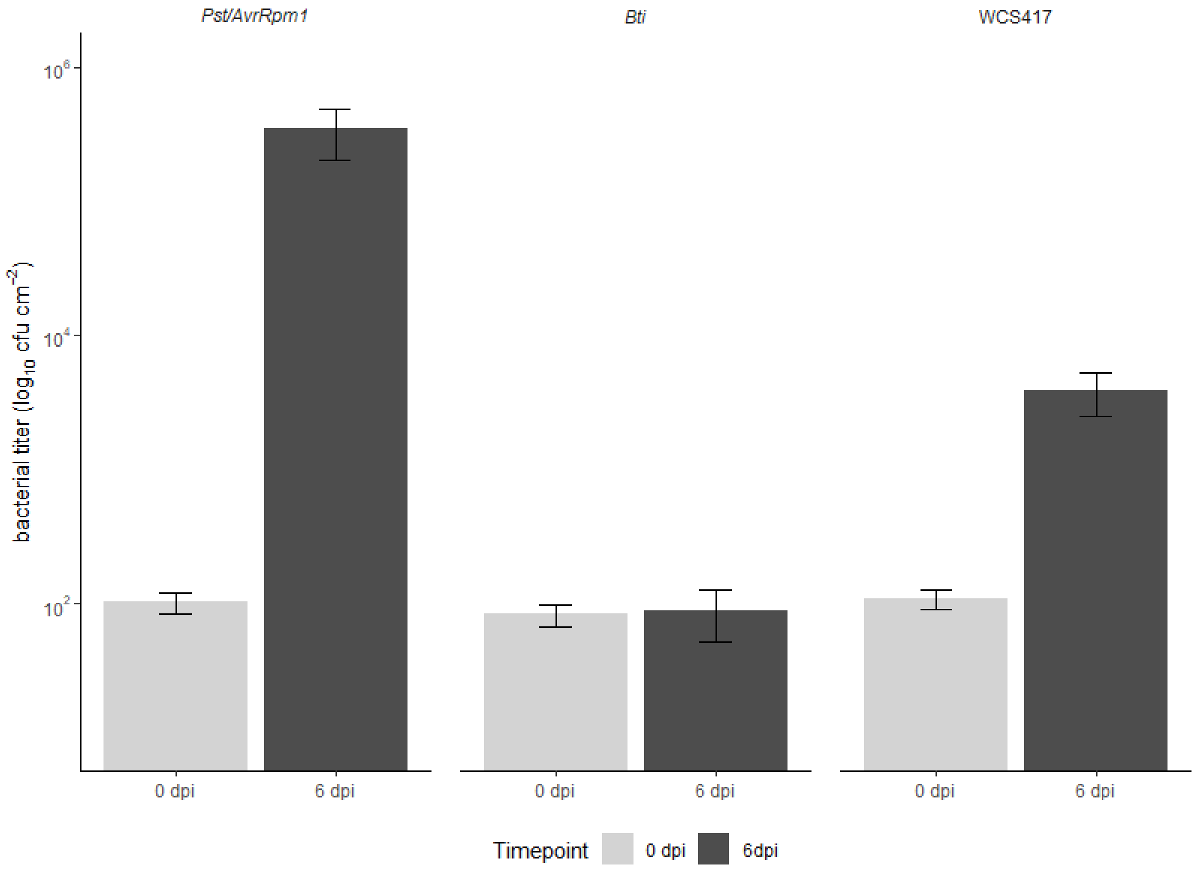


**Supplementary Figure S3**. Growth of ISR-inducing bacteria in *A. thaliana* leaves. Leaves of 4-5-week-old *A. thaliana* plants were syringe-infiltrated with *Pst/AvrRpm1* (positive control), *Bti*, or WCS417 as indicated above the panel. The resulting *in planta* titers of these bacteria were monitored 2 h after inoculation (0 dpi) and six days later (6 dpi). Bars represent the mean of two biologically independent experiments, including three replicates each ± SD.

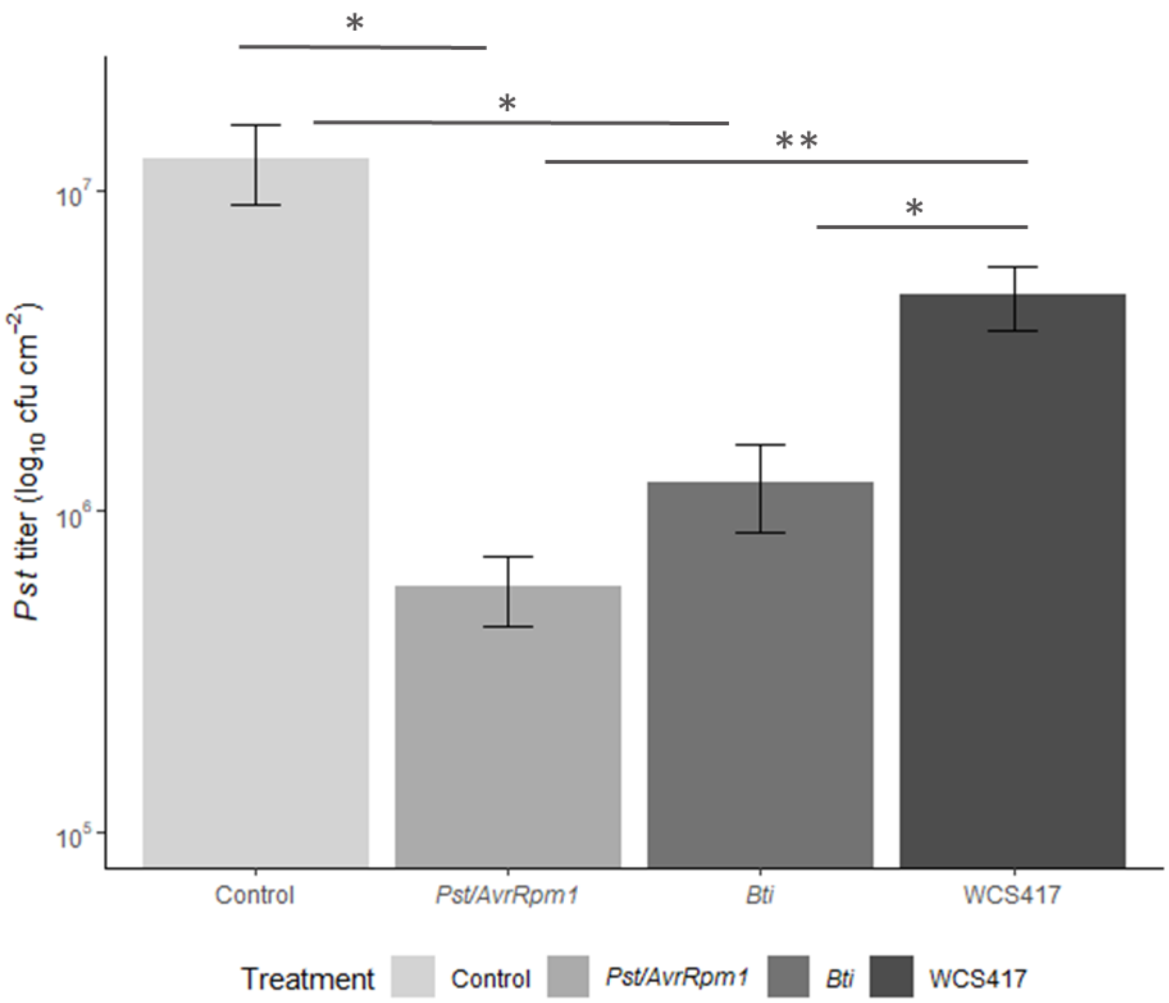


**Supplementary Figure S4**. Systemic immunity in response to local leaf application of *Bti* and WCS417. The first two true leaves of 4–5-week-old *A. thaliana* plants (Col-0) were infiltrated with *Pst* carrying the effector *AvrRpm1* (positive control), *Bti*, WCS417, or a corresponding (negative) control solution (Control). Three days later, two systemic leaves were challenged with *Pst* by syringe infiltration. Bars represent the mean *in planta Pst* titers at 4dpi from three biologically independent experiments, including three replicates each ± SD. Asterisks indicate significant differences between the treatments indicated by the corresponding lines (pairwise Wilcoxon test, adjusted for multiple testing by Benjamini-Hochberg procedure, *, $p$ <0.05, **, $p$ <0.01).

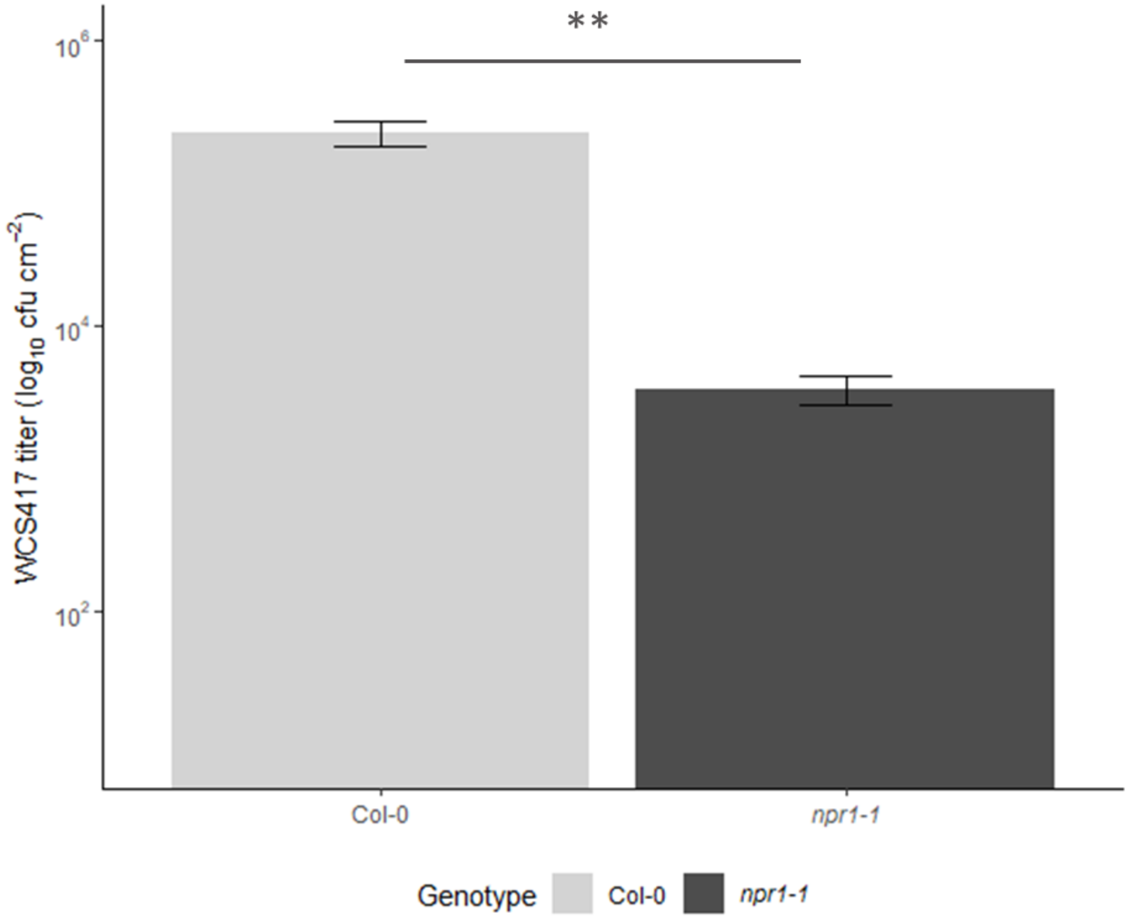


**Supplementary Figure S5**. WCS417 titers in Col-0 (wild type) and *npr1-1* mutant plants. Leaves of 4-5-week-old *A. thaliana* plants of the genotypes indicated below the panel were syringe-infiltrated with WCS417. The resulting *in planta* WCS417 titers are shown at 6 dpi. Bars represent the mean of two biologically independent experiments, including three replicates each ± SD (Student's t-test, *, $p$ <0.05, **, $p$ <0.01).
